# Zetaproteobacteria pan-genome reveals candidate gene cluster for twisted stalk biosynthesis and export

**DOI:** 10.1101/2021.03.12.435129

**Authors:** E. Koeksoy, O.M. Bezuidt, T. Bayer, C.S. Chan, D. Emerson

## Abstract

Twisted stalks are morphologically unique bacterial extracellular organo-metallic structures containing Fe(III) oxyhydroxides that are produced by microaerophilic Fe(II)-oxidizers belonging to the Betaproteobacteria and Zetaproteobacteria. Understanding the underlying genetic and physiological mechanisms of stalk formation is of great interest based on their potential as novel biogenic nanomaterials and their relevance as putative biomarkers for microbial Fe(II) oxidation on ancient Earth. Despite the recognition of these special biominerals for over 150 years, the genetic foundation for the stalk phenotype has remained unresolved. Here we present a candidate gene cluster for the biosynthesis and secretion of the stalk organic matrix that we identified with a trait-based analyses of a pan-genome comprising 16 Zetaproteobacteria isolate genomes. The “**s**talk **f**ormation in **Z**etaproteobacteria” (*sfz*) cluster comprises six genes (*sfz1-sfz6*), of which *sfz1* and *sfz2* were predicted with functions in exopolysaccharide synthesis, regulation, and export, *sfz4* and *sfz6* with functions in cell wall synthesis manipulation and carbohydrate hydrolysis, and *sfz3* and *sfz5* with unknown functions. The stalk-forming Betaproteobacteria *Ferriphaselus* R-1 and OYT-1, as well as dread-forming Zetaproteobacteria *Mariprofundus aestuarium* CP-5 and *Mariprofundus ferrinatatus* CP-8 contain distant *sfz* gene homologues, whereas stalk-less Zetaproteobacteria and Betaproteobacteria lack the entire gene cluster. Our pan-genome analysis further revealed a significant enrichment of clusters of orthologous groups (COGs) across all Zetaproteobacteria isolate genomes that are associated with the regulation of a switch between sessile and motile growth controlled by the intracellular signaling molecule c-di-GMP. Potential interactions between stalk-former unique transcription factor genes, *sfz* genes, and c-di-GMP point towards a c-di-GMP regulated surface attachment function of stalks during sessile growth.

## 1 Introduction

Twisted stalks are organo-mineral composites that were first recognized in the early 19^th^ century for their distinctive ribbon-shaped morphology (Ehrenberg, 1836). They are formed by neutrophilic microaerophilic chemolithoautotrophs that obtain energy from the enzymatic oxidation of Fe(II) to Fe(III) with oxygen (O_2_) as the terminal electron acceptor (Emerson and Moyer, 1997; Emerson and Revsbech, 1994), and that classify into the Betaproteobacteria (Kato et al. 2015, 2013) and Zetaproteobacteria (Emerson et al., 2007). Only four isolates of the Gallionellaceae family within the ecologically and physiologically diverse Betaproteobacteria have been characterized as freshwater Fe(II)-oxidizers to date, with two of them lacking the stalk phenotype (Emerson et al., 2013; Kato et al., 2015). Contrastingly, all previously reported cultured Zetaproteobacteria are characterized as marine Fe(II)-oxidizers (Emerson et al., 2007), with two isolates reported as being additionally capable of H2 oxidation (Mori et al., 2017). The full metabolic potential of the Zetaproteobacteria is yet to be uncovered (Field et al., 2015; Brooks and Field, 2020) and so are the evolutionary grounds for the rise of the stalk formation phenotype in two distinct Proteobacteria classes.

Their requirement for oxygen as terminal electron acceptor limits microaerophilic Fe(II)-oxidizers to low-O_2_ environmental niches (typically <50 μM) where they can outcompete the kinetics of chemical Fe(II) oxidation through ambient O_2_ (Druschel et al., 2008; Rentz et al., 2007; McAllister et al., 2019). Contrary to their specialized metabolic demands, both culture-dependent and culture-independent approaches revealed that these bacteria occupy a diverse range of habitats, including marine hydrothermal vents and submarine volcanoes (Vander Roost, Thorseth, and Dahle, 2017; Emerson et al., 2017), subtropical coastal catchments (Lin et al., 2012), continental margins (Rubin-Blum et al., 2014), coastal bays (Mumford, Adaktylou, and Emerson, 2016), estuarine water columns (Chiu et al., 2017; Garrison et al., 2019), marine sediments (Laufer et al., 2017; Beam et al., 2018), and worm burrow openings in coastal sediments (McAllister et al., 2015; Beam et al., 2020). The source environments of neutrophilic Fe(II)-oxidizing Betaproteobacteria were primarily reported as ferruginous groundwater discharge points (Emerson et al., 2013; Kato et al., 2015), while metagenomic data also indicates their presence in marine environments such as the Arctic Mid-Ocean Ridge, where they co-occur with Zetaproteobacteria (Vander Roost, Thorseth, and Dahle, 2017).

Stalk formation has been recognized as a characteristic morphological trait of Fe(II)-oxidizing bacteria since the 1800s (Ehrenberg, 1836). It has only been relatively recently that successful isolation of Fe(II)-oxidizing Zeta- and Betaproteobacteria has enabled detailed morphological and chemical characterization of their twisted stalk products. For instance, time lapse microscopy of the Zetaproteobacterium *Mariprofundus ferrooxydans* PV-1 revealed single cells excreting multiple nanometer-thin fibers at the cell concavity that elongate at a rate of 2.2 μm h^-1^ as cells grow and oxidize Fe(II). A contemporaneous rotation of the cell was suggested to cause the helical morphology of stalks (Chan et al., 2011). A combination of spectroscopy and microscopy approaches applied on stalks of isolate cultures, and in environmental Fe-mat samples resolved the stalk nanofibrils to consist of acidic polysaccharides with carboxylic functional groups (Chan et al. 2011, 2009). The organic matrix is proposed to adsorb Fe(III) originating from biotic Fe(II) oxidation (Chan et al., 2004), resulting in its encrustation with amorphous Fe(III) oxyhydroxide (FeOOH) during initial stalk growth. As the stalks age, amorphous FeOOH becomes metastable ferrihydrite which is overprinted with the FeOOH lepidocrocite (Chan et al., 2011). Based on these observations, stalks have been suggested to serve as biomarkers for primordial microaerophilic Fe(II) oxidation in the geologic record (Picard et al., 2015; Chan et al., 2016; Dodd et al., 2017, Krepski et al., 2013; Little et al., 2021), that may be of astrobiological interest as well.

Despite substantial progress on the chemical composition and nano-scale morphology of stalks, the underlying genetic machinery of this unique bacterial product remains unresolved. Certain Fe(II)-oxidizing Beta- and Zetaproteobacteria, i.e. *Sideroxydans lithotrophicus* ES-1, *Gallionella capsiferriformans* ES-2, and *Ghiorsea bivora* TAG-1 and SV-108, were reported not to form twisted stalks but to produce amorphous Fe(III) oxyhydroxides instead (Mori et al., 2017; Emerson et al., 2013). Other Zetaproteobacteria isolates, i.e. *Mariprofundus aestuarium* CP-5 and *Mariprofundus ferrinatatus* CP-8, were demonstrated to excrete shorter and thicker exopolymers encrusted in Fe(III) oxyhydroxides that differ morphologically from stalks, known as ‘dread’-biominerals due to their unique morphology (Chiu et al., 2017). Collectively, these observations imply that stalks are not essential for the survival of Fe(II)-oxidizing bacteria. However, since stalk-biosynthesis must incur some bioenergetic cost, it is logical that they confer physiological and/or ecological advantages over Fe(II)-oxidizers that form amorphous Fe(III) oxides. Proposed functions include anchoring of the cell in environments with favorable gradients of Fe(II) and O_2_, and prevention of cell encrustation in Fe(III) through templating its sorption to an organic matrix (Chan et al., 2011).

Identifying the genes that encode stalk formation is a vital step in deciphering the evolutionary and physiological relevance of this unique bacterial trait. The organic nature of the stalk fibrils together with their secretion from the cell surface suggest an interaction of multiple genes with different functions, including carbohydrate synthesis, secretory pathways, and cellular export. Furthermore, we hypothesize that there may be a reciprocal regulation with Fe(II) oxidation pathways which requires special attention in the search for candidate genes involved in stalk biomineralization.

Kato et al. (2015) previously reported a cluster of four genes with putative involvement in stalk formation that are shared between stalk-forming freshwater Fe(II)-oxidizers and the Zetaproteobacteria *M. ferrooxydans* PV-1, M34, and *Mariprofundus* sp. EKF-M39. Of these, three genes were identified to be similar to the exopolysaccharide synthesis encoding *xagBCD* genes of plant-pathogenic *Xanthomonas* species that attach to their targets with a xanthan-based biofilm (Tao et al., 2010; Zhang et al., 2013). The fourth gene was annotated as a BcsB-like cellulose synthase regulator protein (Kato et al., 2015). Due to the lack of a systematic comparison of all currently available Zetaproteobacteria isolate genomes, we chose to search for genes connected to stalk formation with a pan-genome approach and subsequently compared our results to those of Kato et al. (2015). Here, we describe the size and trajectory of a pan-genome that comprises 16 genomes of cultured Zetaproteobacteria isolates. Our data reveals substantial genetic diversity within the Zetaproteobacteria, and provides insight into gene clusters that are likely involved in stalk generation. These results on stalk formation align with, and significantly extend those from Kato et al. (2015), while also pointing towards a potential signaling cascade that may regulate a switch between sessile and motile growth in Zetaproteobacteria.

## 2 Material & Methods

### 2.1 Functional annotation of Zetaproteobacteria genomes

All 16 sequenced genomes of cultured Zetaproteobacteria isolate strains (Table 1) were retrieved from the Joint Genome Institute’s Genome Portal (https://genome.jgi.doe.gov/portal/). Automated functional annotation of all genomes was conducted using Prokka (Seemann, 2014), which predicted coding sequences using Prodigal (Hyatt et al., 2010), ribosomal RNA genes using RNAmmer (Lagesen et al., 2007), transfer RNA genes using Aragorn (Laslett and Canback, 2004), signal leader peptides using SignalP (Petersen et al., 2011), and non-coding RNA regions using Infernal (Nawrocki, Kolbe, and Eddy, 2009). Using BLAST+ (Camacho et al., 2009) and HMMER (Eddy, 2011), Prokka hierarchically annotates against the UniProt (Apweiler et al., 2004), RefSeq (O’Leary et al., 2016), Pfam (Punta et al., 2012), and TIGRFAMs (Haft et al., 2013) databases to predict gene functions for previously identified coding sequences.

**Table 1:**
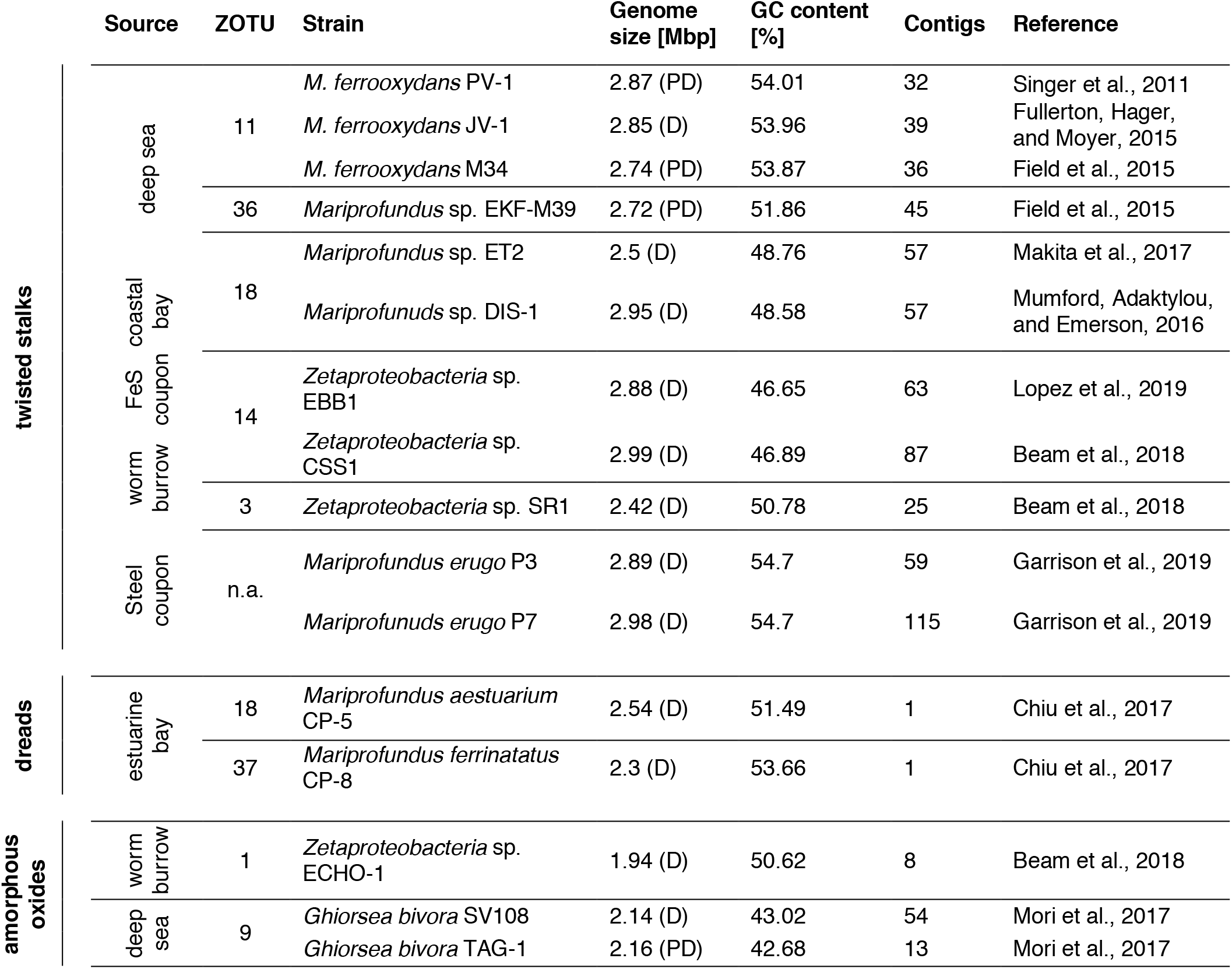
Source environments, biomineral phenotypes, and general genome features of cultured Zetaproteobacteria isolate genomes included in this pan-genome analyses. ZOTU – Zetaproteobacteria operational taxonomic unit (McAllister, Moore & Chan, 2018); PD – permanent draft; D – Draft; n.a. – not available.

### 2.2 Calculation of average amino acid identities among Zetaproteobacteria

We identified the degree of genomic similarity between all Zetaproteobacteria and Betaproteobacteria genomes included in our study by calculating their average amino acid identity (AAI) using CompareM (https://github.com/dparks1134/CompareM) with default parameters of 30% sequence identity and 70% sequence alignment length. We computed Bray-Curtis dissimilarity and average linkage hierarchical clustering using the R package vegan (Oksanen et al., 2016), and visualized our results with the R libraries gplots (Warnes et al., 2016), Heatplus (Ploner, 2019), and RColorBrewer (Neuwirth, 2014).

### 2.3 Pan-genome computation

We used functionally annotated genome files (see section 2.1) of 16 Zetaproteobacteria isolates (Table 1) for computing the size and trajectory of 7 pan-genomes that differed in minimum blastp percentage identities between 30-90% (Altschul et al., 1990) using the Roary software (Page et al., 2015). Briefly, Roary converted coding regions identified by Prokka (Seemann, 2014) into protein sequences, removed partial sequences, and iteratively pre-clustered coding regions with CD-HIT (Fu et al., 2012). After pairwise sequence alignment with blastp (Altschul et al., 1990), Roary clustered sequences with the Markov cluster algorithm (MCL) (Enright, Van Dongen, and Ouzounis, 2002) with an MCL inflation value of 1.5 and subsequently merged pre-clustering results from CD-HIT with those from MCL (Page et al., 2015). We visualized the calculated pan-genome size and trajectory with R using the ggplot2 package (Wickham, Chang, and Wickham, 2016).

### 2.4 Functional analyses of pan-genome subdivisions and individual genomes

We analyzed the pan-genomes in three categories as defined by Koonin and Wolf (2008), i.e. (1) core (genes present in all 16 genomes), (2) accessory (genes present in 2-15 genomes), and (3) strain-specific genes (genes present in only 1 genome). After extracting core, shell, and unique genes from the pan-genome computed at 50% minimal blastp identity using the ‘query_pan_genome −a *.gff’ script of Roary (Page et al., 2015), we analyzed the distribution and enrichment of clusters of orthologous groups (COGs) in each pan-genome subdivision and in individual genomes by assigning COG categories to query protein sequences using cdd2cog (Leimbach, 2016).

### 2.5 Identification of stalk-associated genes

Using the ‘query_pan_genome −a difference’ function in Roary (Page et al., 2015) on previously computed pan-genomes, we extracted genes uniquely shared among all 11 stalk-former genomes (Table 1), that at the same time were absent in the genomes of stalk-less isolates. Protein coding sequences that were initially annotated as hypothetical proteins by Prokka (Seemann, 2014) were further analyzed for functional information using EggNOG mapper (Huerta-Cepas et al., 2019) and InterProScan (Quevillon et al., 2005). We computed multiple sequence alignments with protein sequences of potential candidate genes for stalk formation with Muscle (Edgar, 2004) and visualized our results with Jalview2 (Waterhouse et al., 2009) to search for highly conserved regions and identical amino acid residues. Using Gene Graphics (Harrison et al., 2018), we generated synteny plots of putative stalk formation genes in the genomes they were found in, given that they were available on the National Center for Biotechnology Information (NCBI) database.

### 2.6 SEM sample preparation and imaging of cell-stalk-aggregates

We obtained an active culture of *Mariprofundus micogutta* ET2 from the Japan Collection of Microbes (Ibaraki, Japan) that we cultivated on zero-valent iron plates with artificial seawater medium at 5% headspace oxygen after Laufer et al. (2016) for 5 to 8 days prior to analyses with an Olympus BX60 epifluorescence microscope (Olympus, Japan) and a Qicam 1394 (Qimaging, Canada). Similarly, we cultivated *Mariprofundus* sp. DIS-1 on zero-valent iron plates with artificial seawater medium after Laufer et al. (2016) for 5 days prior to harvesting 1 mL culture for sample preparation.

To ensure good preservation of biological and mineralogical structures, samples were split up for different treatments: either glutaraldehyde fixation followed by stepwise dehydration or air-drying. Biological samples were fixed in 2.5% glutaraldehyde on ice for 3 hours. Samples were then washed three times with ultrapure water, pelleted at 2,800 g for 1 minute, and mounted on poly-L-lysine (0.1% w/v aqueous solution) coated glass slides. Samples were then dehydrated sequentially with 30%, 50%, 70%, and 95% ethanol for 5 minutes each, followed by two times 100% ethanol for 30 minutes each. Afterwards, samples were treated twice with hexamethyldisilazan for 30 seconds each. After final air drying, samples were stored in a dry chamber at room temperature.

Non-fixed samples for preservation of mineral structures were washed three times with ultrapure water, concentrated by centrifugation at 2,800 g for 1 minute, mounted on poly-L-lysine glass slides, and air-dried afterwards. All sample-slides were applied onto carbon-taped aluminum stubs and sputter-coated for 120 seconds at 4×10^-2^ bar to achieve a 12 nm platinum layer using a Bal-Tec SCD005 sputter coater (Baltic praeparation, Germany). Imaging was performed with a JSM-6500F field emission scanning electron microscope (JEOL, Germany) with a Schottky field emitter at 5kV acceleration voltage and a working distance of approximately 10 mm at the center for Light-Matter Interaction, Sensors and Analytics (LISA+, University of Tuebingen, Germany).

## 3 Results & Discussion

The diverse range of geographically distinct Fe(II)-rich environments inhabited by Zetaproteobacteria, including terrestrial, freshwater, coastal, and deep-sea environments (Table 1), hints towards their adaptive nature. Although the number of isolates is relatively limited across all these environments, there was no obvious differentiation of morphological type to a specific habitat-type (Table 1). We refer to the dread and amorphous oxide phenotypes collectively as ‘stalk-less’ hereafter.

### 3.1 Pan-genome reveals high genetic diversity among Zetaproteobacteria

The pan-genome is defined as the entire repertoire of genes accessible to a bacterial clade, subdivided into conserved ‘core’ genes (shared between all *n* genomes), variable ‘accessory’ genes (shared between two and *n*-1 genomes), and ‘strain-specific’ genes (present in only one genome) (Koonin and Wolf, 2008; Vernikos et al., 2015; Tettelin et al., 2005). The relative proportions of these three subdivisions characterize a clade’s genetic and functional richness and its capacity to acquire exogeneous DNA, and define the trajectory of its pan-genome (Rouli et al., 2015). Roary regression curve models predicted the Zetaproteobacteria pan-genome to be open with infinite growth in total gene numbers and decreasing core gene numbers as new genomes are added (Figure 1). Accessory and strain-specific genes made up the majority of the Zetaproteobacteria pan-genome, whereas the core genome was relatively small with only 0.04-10.3% of the total gene count (90-30% blastp identity, respectively; Table S1).

**Figure 1:**
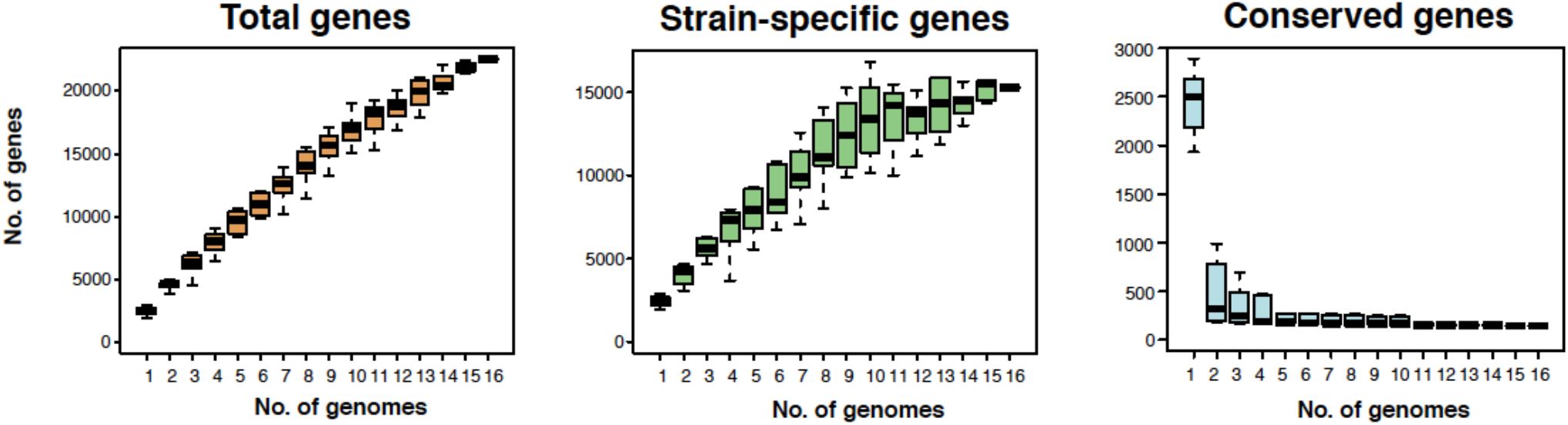
Size and trajectory of the Zetaproteobacteria pan-genome with the number of total, strain-specific, and conserved genes as genomes were added up to a total of 16. Data is shown for a pan-genome computed at a minimal blastp identity of 80%.

Open pan-genomes generally reflect sympatric lifestyles, i.e. life in mixed communities with high rates of horizontal gene transfer (Rouli et al., 2015; Vernikos et al., 2015), and are associated with niche versatility (Tettelin et al., 2005). Our results are consistent with the previously reported high genomic diversity across Zetaproteobacteria based on metagenomic data (Fullerton et al., 2017; McAllister et al., 2019) and indicate that Zetaproteobacteria may undergo considerable genetic exchange with other community members. We could not identify a correlation between source environment and the proportion of strain-specific and shared genes in isolate genomes (Table S2) and so cannot conclude that a specific habitat may favor higher gene acquisition rates in Zetaproteobacteria. The high genomic diversity among currently available Zetaproteobacteria isolates is further reflected in amino acid identities (Figure S1) that classify only deep sea members *M. ferrooxydans* PV-1, JV-1, and M34 (98.1±5.4% to 99.9±1.0%), and coastal estuarine isolates *M. erugo* P3 and P7 (98.9±5.0%; Figure S1) at the species rank (AAI of >95%; Medlar, Törönen, and Holm, 2018) as previously reported by McAllister et al. (2019, 2020) and Garrison et al. (2019).

### 3.2 Candidate gene cluster for stalk synthesis and export is shared with stalk-forming Betaproteobacteria

We used comparative genomics as an approach to identify the genetic basis for stalk formation where each Zetaproteobacteria isolate genome was assigned to either the stalk-former, dread-former, or amorphous-oxide-former trait groups (Table 1). While the assignment was clear for most isolate genomes from the documentation in the literature (Mori et al., 2017; Chiu et al., 2017; Mumford, Adaktylou, and Emerson, 2016; Chan et al., 2011; Beam et al., 2018; Field et al., 2015) or from microscopic observations in our laboratory (Figure 2A-F), the trait was more ambiguous for strain *Mariprofundus micogutta* ET2 (JCM 30585^T^), a member of the Zetaproteobacteria operational taxonomy unit 18 (ZOTU 18). Makita et al. (2017) reported strain ET2 to form thin Fe- and C-rich extracellular filaments that differ morphologically from stalks. Using light microscopy, we identified stalk production in an active culture of *M. micogutta* ET2 (Figure 2G-H), although the majority of cells did not appear associated with obvious stalk structures. Hence, we assigned ET2 to the ‘stalk-forming’ trait group that overall comprised 11 genomes (PV-1, JV-1, M34, ET2, DIS-1, EKF-M39, EBB1, CSS1, SR1, P3, and P7), while we assigned 5 genomes to the ‘stalk-less’ (CP5, CP8, TAG-1, SV-108, and ECHO1) trait group during our analyses.

**Figure 2:**
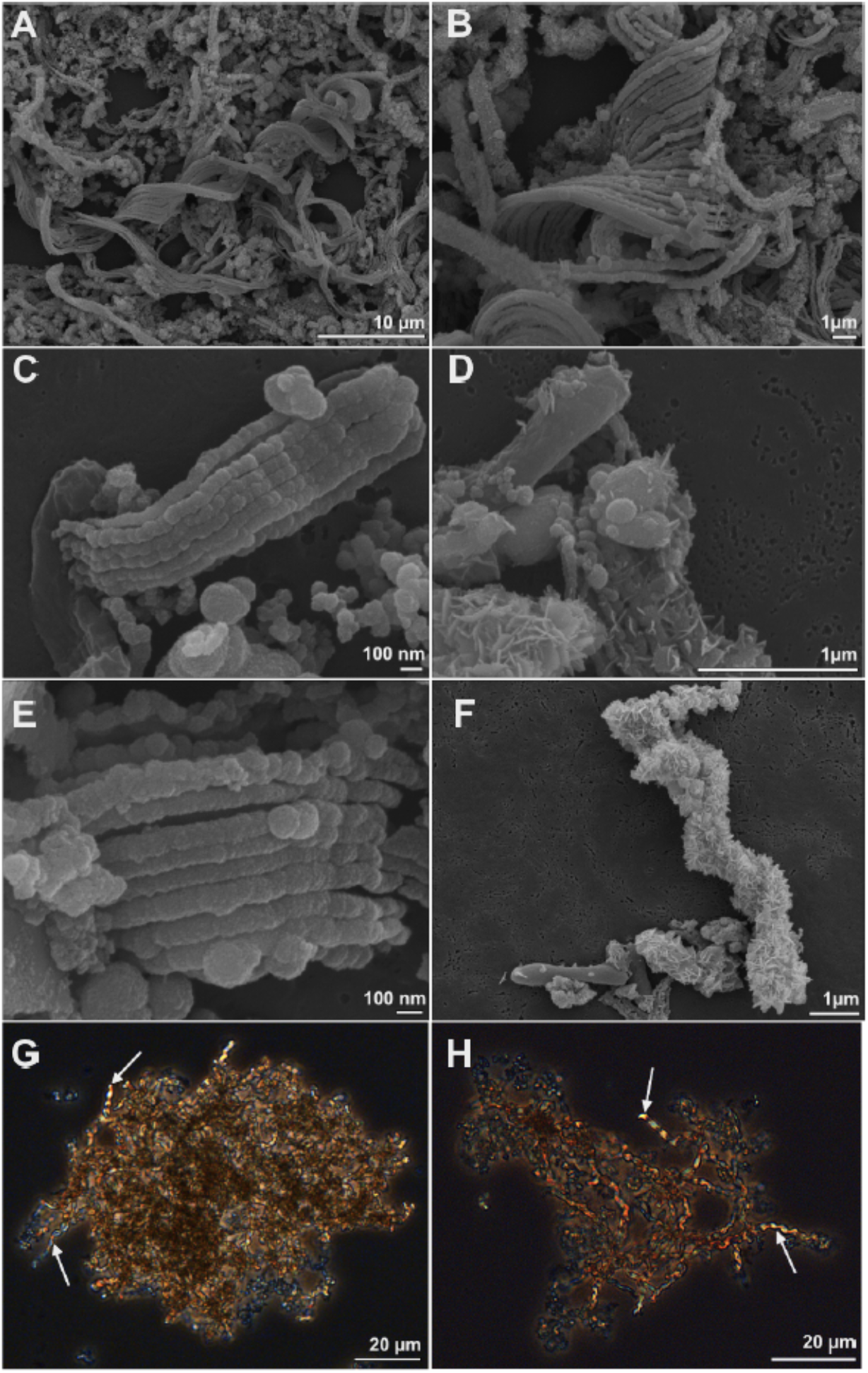
Scanning electron microscopy (A-F) and light microscopy images (G-H) of cell-stalks aggregates in cultures of *Mariprofundus* sp. DIS-1 (A-F) and *M. micogutta* ET2 (G-H). White arrows in G-H indicate stalks.

We extracted all genes from the Zetaproteobacteria pan-genome that are shared among stalk-former genomes and at the same time are lacking in stalk-less Zetaproteobacteria during our analysis. Our search revealed 82 genes unique to stalk-forming Zetaproteobacteria at minimal blastp identities between 30-80% (Table S3). We excluded genes that were identified above 90% minimal blastp identity from our analysis as they were only shared between 8 or less stalk-formers. Prokka annotated 34 stalk-former unique genes with functions in flagellar basal-body biosynthesis, motility, chemotaxis, and terminal electron transfer with cbb3-type cytochromes (Table S3; see sections 3.7 and 3.8 for further details). The other 48 genes were predicted as hypothetical proteins among which we found 6 conserved genes that cluster together with a shared synteny in all stalk-former genomes.

The high level of conservation above 50% amino acid identity (Table 2) and its lack in stalk-less Zetaproteobacteria emphasize the potential for this gene cluster as a candidate for the biosynthesis and export of the stalk organic matrix, which we therefore refer to as the **s**talk **f**ormation in **Z**etaproteobacteria (*s/z*) cluster with genes *sfz1-sfz6* hereafter. Protein motif analysis with InterProScan (Quevillon et al., 2005) predicted domains with functions in exopolysaccharide synthesis and export, cell wall synthesis inhibition, and carbohydrate hydrolysis in Sfz1, Sfz2, and Sfz4, respectively, that we describe in detail below (see sections 3.3 and 3.4). We could not identify known domains and putative functions for Sfz3, Sfz5, and Sfz6 (see section 3.5).

**Table 2:** Sfz genes with minimal and maximal basepair length in different stalk-former genomes at minimal blastp identities between 50-70%. EPS=exopolysaccharide.

| **Gene** | **Annotation** | **Putative function** | **Min.**<br>**length** | **Max.**<br>**length** | **Min.**<br>**blastp**<br>**identity** |
| --- | --- | --- | --- | --- | --- |
| *sfz1* | hypothetical protein | Regulator of EPS synthase | 1067 | 2576 | 60 |
| *sfz2* | hypothetical protein | EPS synthesis and export | 2051 | 2084 | 70 |
| *sfz3* | hypothetical protein | unknown | 1499 | 1535 | 60 |
| *sfz4* | hypothetical protein | carbohydrate/ cell wall hydrolysis | 971 | 1037 | 50 |
| *sfz5* | hypothetical protein | unknown | 929 | 983 | 50 |
| *sfz6* | hypothetical protein | unknown | 3311 | 7352 | 50 |

The appearance of twisted stalks in Zeta- and Betaproteobacteria indicates that this trait evolved from shared ancestral genes encoding stalk biosynthesis and export. We elucidated the presence and degree of conservation of the *sfz* cluster in stalk-forming Betaproteobacteria using blastp (Altschul et al., 1990), and found homologs of *sfz1-sfz4* with a shared synteny in the stalk-formers *Ferriphaselus amnicola* OYT-1 and *Ferriphaselus* sp. R-1 (Figure 3; blastp identities below 40%). We refer to *sfz* homologs in Betaproteobacteria as *sfb1-sfb4* (**s**talk **f**ormation in **B**etaproteobacteria) hereafter. As for the *sfz* cluster in Zetaproteobacteria, we found no *sfb* genes in the stalk-less Fe(II)-oxidizing Betaproteobacteria *Gallionella capsiferriformans* ES-2 and *Sideroxydans lithotrophicus* ES-1. Our blastp search further revealed the dread-forming Zetaproteobacteria *M. aestuarium* CP-5 and *M. ferrinatatus* CP-8 to comprise less conserved homologs of *sfz1-sfz6* (blastp identities below 40%, Figure 3), which may point towards two distinct phenotypes among Zetaproteobacteria that are rooted in the same gene cluster.

**Figure 3:**
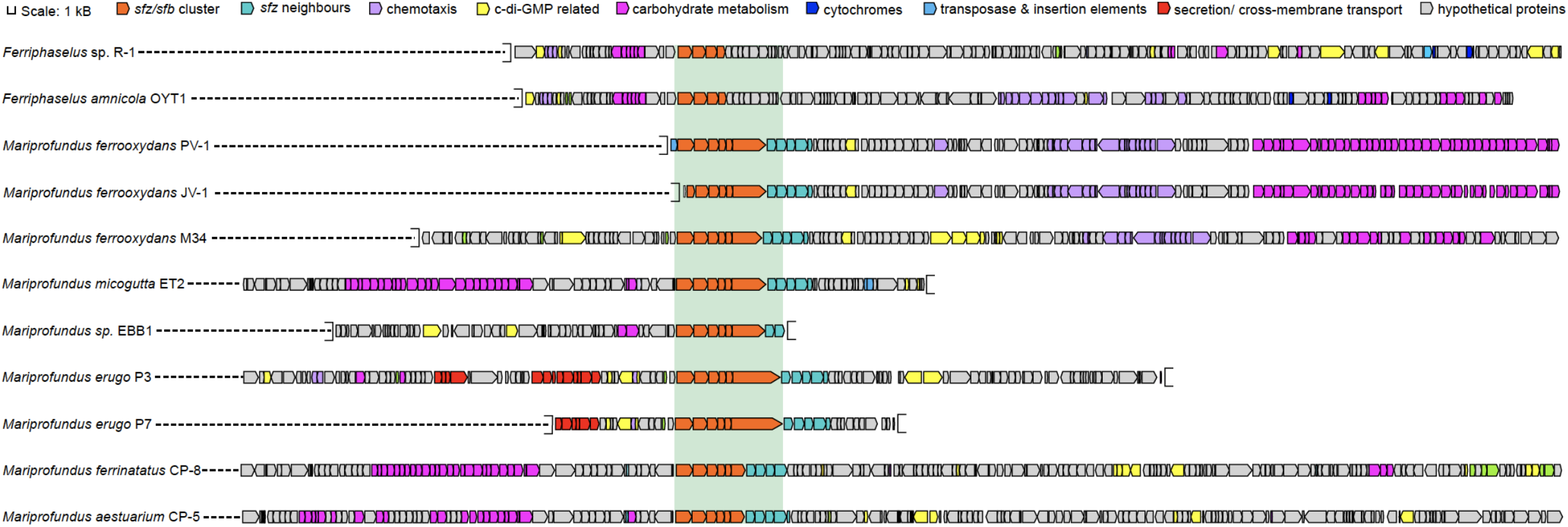
Gene synteny of the *sfz/sfb* cluster in stalk-forming Zetaproteobacteria and Betaproteobacteria. *Sfz* clusters are only shown for genomes that are already publicly available.

Our findings align with a previous report of the *sfb* and *sfz* genes by Kato et al. (2015) as the ‘*xag*-like’ gene cassette (CDS9-CDS12), that they found to be shared between freshwater stalk-formers and marine *M. ferrooxydans* PV-1, M34, and *Mariprofundus* sp. EKF-M39. Based on this limited genomic data, the authors surmised that respective genes might be involved in stalk formation. Our results here significantly expand the number of genomes, especially among the Zetaproteobacteria, and provide additional comparative analysis between stalk-forming and stalk-less strains. Thus, our present data support the proposal of Kato et al. and underline the strong potential of this gene cassette as a candidate gene cluster for stalk exopolysaccharide synthesis. Interestingly, Betaproteobacteria lack homologues for *sfz5* and *sfz6,* indicating that only *sfz1-sfz4* and *sfb1-sfb4* may be essential for the stalk phenotype, whereas the roles for *sfz5* and *sfz6* in stalk-formation may be more ambiguous. In general, exopolysaccharide synthesis strategies can vary significantly among bacteria, and are encoded by different sets of genes that can be more or less conserved in sequence and operon structure across taxa (for a review see Schmid et al., 2015). Thus, it is possible that additional accessory genes play a structural role in stalk-formation in the strains they are found in, but that other strains may use different structural polysaccharides in stalk-formation.

### 3.3 Sfz1 is a putative c-di-GMP-dependent regulator of exopolysaccharide synthesis

Our protein motif analyses predicted the Sfz1 amino acid sequence (60% min. blastp identity) to contain two non-cytoplasmic BcsB-like domains flanked by an N-terminal transmembrane signal peptide domain and a C-terminal transmembrane domain. We found the N-terminal part of Sfz1 to be less conserved than the C-terminal fraction that contains 76 identical residues in all stalk-formers (Figure S2). *M. ferrooxydans* JV-1 only contains the second part of *sfz1*, which likely is an artifact as the gene is split over two contigs in its genome.

BcsB is the periplasmic membrane-anchored regulatory subunit of the dimeric cellulose synthase enzyme in *Acetobacter xylinum* (Wong et al., 1990) that regulates the polymerization of uridine 5’-disphosphate glucose (UDP-glucose) to cellulose, an exopolysaccharide that builds the structural basis of a variety of biofilms. The reaction is catalyzed by the family II glycosyltransferase BcsA upon activation by the BcsB subunit, which depends on the binding of the positive effector cyclic dimeric guanylate monophosphate (c-di-GMP). High intracellular c-di-GMP levels were shown to stimulate exopolysaccharide synthesis and biofilm formation as part of surface attachment and sessile growth in different bacteria, including *Pseudomonas aeruginosa, Escherichia coli, Salmonella enterica serovar Typhimurium,* and *Shewanella oneidensis* MR-1. In contrast, low c-di-GMP levels induce cell detachment and motility (Kuchma et al., 2015; Merritt et al., 2007; Simm et al., 2004).

Generally, c-di-GMP was found to be a central signaling molecule for a variety of signaling cascades, and it particularly controls the switch between motile and sessile growth in bacteria, where biofilm formation is up- or downregulated for surface attachment or detachment (for a review see Valentini and Filloux, 2016; D’Argenio and Miller, 2004). An interaction between Sfz1 and c-di-GMP could point towards a surface holdfast function of stalks as previously proposed by Singer et al. (2011), and our pan-genome data provides additional hints for such a functional relationship (see section 3.7). Our detailed analysis of Sfz1 indicates it could encode the conserved c-di-GMP binding motifs RxxD (glutamine-x-x-aspartate; Figure S2, residues 295-298 and 830-833; Chou & Galperin, 2016) and RxxxR (glutamine-x-x-x-glutamine; Figure S2, residues 227-231 and 326-330, Morgan et al., 2014). Sfb1 similarly contained c-di-GMP binding sites with an additional RxxD motif compared to Sfz1 (Figure S2, residues 270-273), whereas it lacked the C-terminal RxxD motif in Sfz1 (residues 830-833) and contained aspartic acid instead of arginine at the RxxxR site at residues 326-330.

Functional exopolysaccharide synthesis would require at least a second gene that encodes a glycosyltransferase function for polysaccharide polymerization similar to BcsA, which we identified to be the case for *sfz2* (see section 3.4). Interestingly, the BcsA and BcsB couple was previously proposed to carry out both the synthesis and translocation of exopolysaccharides, as BcsA can, in addition to its glycosyltransferase activity, also form a channel across the cell membrane that allows for the export of the synthesized polysaccharide chain (Omadjela et al., 2013; Morgan, Strumillo, and Zimmer, 2013). Whether Sfz1 and Sfz2 can carry out a similar function in stalk-formers remains to be explored.

### 3.4 Sfz2 putatively catalyzes exopolysaccharide synthesis and export

Our analyses predicted the highly conserved Sfz2 sequence (70% minimal blastp identity; Figure S3) to contain an N-terminal EpsE-like type II secretion system (T2SS) protein domain (IPR007831) and a C-terminal family II glycosyltransferase domain (IPR001173). The presence of the conserved glycosyltransferase motifs QxxRW (glutamine-x-x-arginine-tryptophane; residues 523-527), DxD (aspartate-x-aspartate; residues 488-490), and TED (threonine-glutamate-aspartate; residues 486-488; Figure S3; Morgan et al., 2016) supports a glycosyltransferase function at the C-terminal end of Sfz2. We found the QxxRW and DxD motifs in Sfb2 as well, whereas it lacked the TED site.

The N-terminal part of Sfz2 may involve a translocation function, as EpsE is a cytosolic, hexameric ATPase that causes asymmetric conformational rearrangements in the T2SS inner membrane platform through ATP hydrolysis, promoting the extension of secreted exopolymers (Korotkov et al., 2012). Guttenplan, Blair, and Kearns (2010) proposed EpsE in *B. subtilis* to be a bifunctional enzyme where it acts either as a family II glycosyltransferase (Garinot-Schneider et al., 2000), or as a molecular clutch that controls the flagellar rotor and inhibits cellular motility (Blair et al., 2008). Due to its gene architecture Sfz2 may encode a similar function, although a translocation function seems questionable. Functional exopolymer secretion via T2SS was demonstrated to require at least 12 different proteins that are generally encoded on a single operon (Sandkvist, 2001) and no other *eps* homologues were identified in the *sfz* cluster or its gene neighborhood (see section 3.6).

Importantly, *eps* gene expression was demonstrated to be upregulated by c-di-GMP in *Vibrio cholerae* (Sloup et al., 2017). As in Sfz1, we found the c-di-GMP binding motifs RxxxR (Figure S2, residues 522-526, Morgan et al., 2014) and RxxD (Figure S2, residues 97-100 and 325-328) in Sfz2 and two additional RxxD motifs in Sfb2 (residues 282-285 and 297-300), which further emphasizes a possible connection between the *sfz and sfb* gene clusters and a c-di-GMP regulated switch into sessile or motile growth in stalk-formers (see section 3.7 for further details). Together, the conservation within Sfz2 with these different motifs involved in exopolysaccharide production and environmental response is circumstantial evidence for similar functions in stalk formation; however experimental work will be necessary to confirm these functions.

### 3.5 Sfz4 and Sfz6 contain domains for carbohydrate cleavage and cell wall synthesis inhibition

The 323-345 aa long Sfz4 sequence (50% minimal blastp identity) is predicted with a function as a family 10 glycosyl hydrolase. Glycosyl hydrolases cleave glycosidic bonds between carbohydrates, and family 10 enzymes have been specifically reported with exopolysaccharide-degrading activities as for instance in xylanase, endo-1,3-beta-xylanase, and cellobiohydrolase enzymes (Do et al., 2013; Henrissat, 1991). Cell wall degrading glycosyl hydrolases were demonstrated with roles in the assembly of supramolecular membrane-spanning structures such as secretion systems, and phenotypes of cells with non-functional cell wall hydrolases included impaired biofilm formation, surface attachment, or chemotaxis (Vollmer et al., 2008; Nambu et al., 1999; Mercier et al., 2002; Vermassen et al., 2019). A possible role of Sfz4 in stalk formation could be in hydrolyzing the stalk exopolysaccharide for cell detachment as part of a transition from sessile to motile growth, given that stalks function as surface holdfasts.

*Sfz6* forms an exception to the other *sfz* genes in that it is distantly located from the *sfz* cluster in Zetaproteobacteria sp. SR-1 and EKF-M39. Since stalk-forming Betaproteobacteria lack a homologue for *sfz6*, it is presumably not essential for stalk formation but rather functions as an accessory gene complementing essential stalk formation genes. The 1415-2270 aa long Sfz6 protein sequence (50% blastp identity) contains an N-terminal SH3 (sarcoma homology-3) domain and a C-terminal CotH domain. Among a variety of different cellular functions, SH3 domains were suggested to function as cell wall binding sites during cell wall hydrolysis (Vermassen et al., 2019). In contrast, the function of CotH is presumably determined by additional adjacent protein domains. For instance, cellulose-binding domains may indicate a role of CotH in the formation of the cellulose-degrading multi-enzyme cellulosome complex or generic interactions with the cell wall (Nguyen et al., 2016). Generally, a domain architecture with a combination of SH3 and CotH seems unique and due to the wide variety of functions encoded by SH3 domains, we cannot confer a potential function for *sfz6.*

### 3.6 Sfz gene neighborhood

A cassette of five genes that is directly adjacent to the *sfz* cluster is conserved across stalk-forming Zetaproteobacteria. Their annotated functions are an ATP-ADP antiporter, a phosphomannose-isomerase, a N-acetylmuramoyl-L-alanine amidase, a MsbA-like ABC-type protein exporter, and a protein of unknown function (Figure 3), respectively. Phosphomannose-isomerases interconvert mannose-6-phosphate and fructose-6-phosphate, and were demonstrated to be involved in the synthesis of the bacterial exopolysaccharides xanthane and alginate (Köplin et al., 1992; Shinabarger et al., 1991), whereas N-acetylmuramoyl-L-alanine amidases cleave amide bonds in the net-like peptidoglycan structure and degrade it (Vermassen et al., 2019; Wilmes et al., 2017). None of the five genes are unique to stalk-formers, and Zetaproteobacteria sp. CSS1 and EBB1 contain only the phosphomannose-isomerase and the ATP-ADP antiporter genes in their *sfz* gene cluster neighborhood. Dread-formers contain four of the five conserved genes in the *sfz* gene neighborhood as well, whereas stalk-forming Betaproteobacteria lack the entire cassette. Altogether, their functional connection to carbohydrate metabolism and close association with the *sfz* cluster point towards a potential involvement in stalk formation, possibly as complementary genes.

Generally, our *sfz/sfb* gene neighborhood analysis revealed these gene cassettes could confer functions in carbohydrate metabolism, chemotaxis, extracellular export, electron transfer, and c-di-GMP related signaling. There is variation in the gene architecture of these cassettes across all stalk-formers (Figure 3). It is possible that this diversity could reflect species-specific adjustments to stalk formation and functional roles that are niche-driven; however, the functional assignments are deduced from homologous genes in other bacteria, and the true phenotypic ramifications for different stalk-formers will require further elucidation.

### 3.7 Stalk-former unique genes emphasize a functional connection to c-di-GMP signaling

Zetaproteobacteria are under constant pressure to compete against abiotic Fe(II) oxidation by oxygen (Druschel et al., 2008; Rentz et al., 2007). Therefore, sensing and quickly responding to geochemical changes as well as sustaining stable growth in environments with optimal Fe(II) and O_2_ gradients are essential attributes for their survival. Our pan-genome analysis revealed a general enrichment of COG categories associated with c-di-GMP signaling (CheY-like response regulator domains, GGDEF, AAA-type ATPase, HD-GYP, and PAS/PAC domains, Table S4) across stalk-forming and stalk-less Zetaproteobacteria. The signaling molecule c-di-GMP upregulates biofilm formation and sessile growth at high intracellular concentrations, whereas declining c-di-GMP concentrations result in increasing surface detachment and motile growth (Figure 4). While these results point towards a pronounced importance of switching between motile and sessile growth in stalk-forming and stalk-less Zetaproteobacteria (Figure 4), our domain analysis of *sfz1-6* and additional stalk-former unique genes highlights a particular emphasis of such a switching response in stalk-formers.

**Figure 4:**
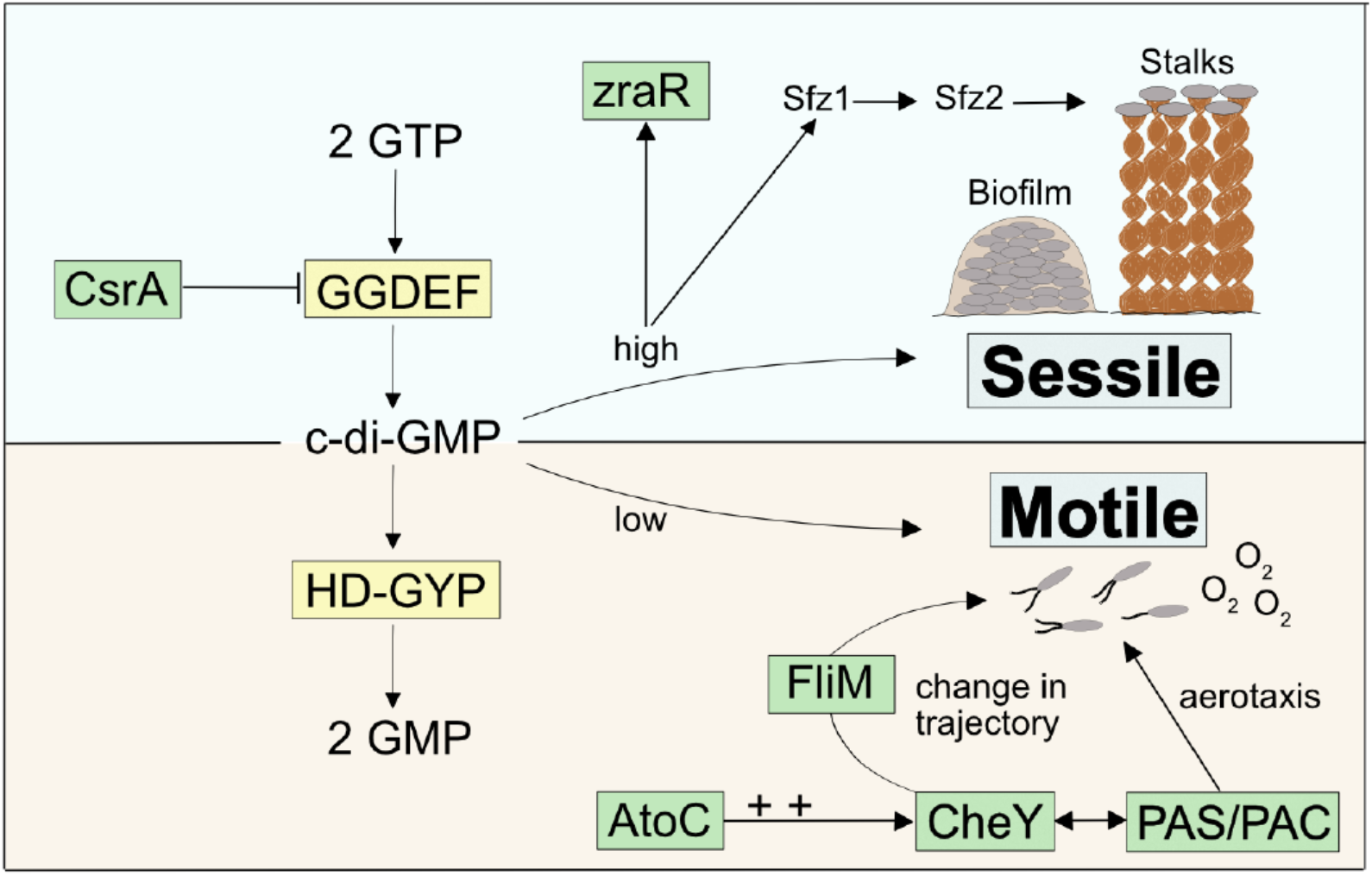
Model for a c-di-GMP regulated signaling cascade in Zetaproteobacteria and potential connections to stalkformer unique genes.

For instance, we found two highly conserved transcriptional regulators unique to stalk-formers that both neighbor additional stalk-former unique genes (Figure S4) and that favor cellular motility, i.e. *atoC* and *csrA* (Table S3; NCBI accession EAU54699.1 and EAU54987.1 in *M. ferrooxydans* PV-1, respectively). Both genes are located in the close neighborhood of c-di-GMP signaling genes with GGDEF- and HD-GYP domains that are known to antagonistically regulate intracellular levels of c-di-GMP (Figure 4). GGDEF-containing diguanylate cyclases catalyze c-di-GMP synthesis and hence an increase in its concentrations (Tal et al., 1998), while HD-GYP-containing phosphodiesterase enzymes catalyze c-di-GMP degradation (Ryan et al., 2009).

CsrA was reported to negatively regulate the expression of GGDEF-containing proteins that upregulate intracellular c-di-GMP levels (Jonas et al., 2008), thereby inhibiting the expression of the exopolysaccharide gene *pgaA* in *Campylobacter jejuni* (Huertas-Rosales et al., 2017). *CsrA-*-mutant cells were demonstrated to show poor swarming ability and lowered resistance to oxidative stress (Fields and Thompson, 2008). AtoC is a transcriptional regulator that was demonstrated to induce the expression of the flagellar *flhDC* and *fliAZY* operons in *E. coli.* Cells deficient in *atoC* were reported to be unresponsive to chemoattractants and showed a general lack of cellular motility. AtoC-dependent regulation of motility and chemotaxis also required an upregulation of CheY (Theodorou et al., 2012), a cytoplasmic chemotaxis response regulator that receives environmental signals and controls cellular swarming trajectories upon phosphorylation. By binding to the flagellar motor protein FliM (Bray and Bourret, 1995; Clegg and Koshland Jr., 1984), phosphorylated CheY induces a clockwise rotation of the flagellum and causes a decrease in torque and swarming speed. In contrast, de-phosphorylated CheY does not bind FliM, which maintains the cellular swarming direction and speed with a counterclockwise flagellar rotation (Figure 4; Scharf et al., 1998; Yuan et al., 2010; Nesper et al., 2017). We found that the stalk-former unique genes comprise the flagellar motor protein gene *fliM* (NCBI accession EAU54562.1 for *M. ferrooxydans* PV-1) within a gene cluster containing other stalk-former unique genes annotated with functions in flagellar motility (Table S3, Figure S4), which may be related to a specialized switching response of the flagellum in stalk-formers, and could indicate a functional connection between stalk formation and chemotaxis regulation.

A third stalk-former unique gene located in the *atoC* gene neighborhood (Figure S4), i.e. *zraR,* (Table S3; NCBI accession EAU54701.1 in *M. ferrooxydans* PV-1), encodes a σ^54^ transcriptional regulator for metal tolerance and resistance (Sallai and Tucker, 2005). Its gene expression was shown to be induced by high intracellular levels of c-di-GMP, although the effect of higher *zraR* transcription has not been resolved yet (Mendez-Ortiz et al., 2006).

### 3.8 Stalk-former unique cytochromes specific to stalk-forming Zetaproteobacteria

In addition to genes with potential functional links to c-di-GMP regulation, we found six stalk-former unique genes with annotations as cbb3-type cytochrome subunits and as *fixGH* genes (Table S3; NCBI accession EAU53376 - EAU53382 in *M. ferrooxydans* PV-1, respectively). The latter are putatively required for the assembly of cbb3-type cytochromes (Preisig et al.,1996) and cluster together in stalk-formers (Figure S4).

Cbb3-type cytochromes are terminal oxidases for aerobic respiration with a high affinity for oxygen (Li et al., 2014; Pitcher and Watmough, 2004) that were reported of being primarily expressed under O_2_-limited conditions (Mandon et al., 1994). They were found among the most highly expressed proteins in the proteome of *M. ferrooxydans* PV-1 (Barco et al., 2015), which is reasonable with respect to the microaerophilic metabolism of Zetaproteobacteria (Field et al., 2015). All Zetaproteobacteria contain cbb3-type cytochrome c genes in their genomes, and the stalk-former unique subunits may be an adaptation required for sensing optimal oxygen levels as part of microaerobic growth during stalk formation.

Generally, Per-Arnt-Sim motifs (PAS) and PAS-associated C-terminal motifs (PAC) were prevalent domains among the strain-specific genes of stalk-forming and stalk-less Zetaproteobacteria (Table S4). PAS/PAC domains are known to respond to environmental signals such as oxygen concentrations, light, and redox potential (Taylor and Zhulin, 1999). The PAC motif was demonstrated to control flagellar rotation during aerotaxis of *E. coli* upon sensing an emitted signal by the PAS domain depending on the surrounding redox conditions (Rebbapragada et al., 1997) with an involvement of CheY (Bibikov et al., 2000). Given their abundance, PAS/PAC domains may determine the swarming trajectories of Zetaproteobacteria during aero- and/or chemotaxis by a similar mechanism (Figure 4).

## 4 Conclusions

Our pan-genome analysis revealed a high genomic diversity among currently available Zetaproteobacteria isolates, and a highly conserved gene cluster in stalk-forming Zetaproteobacteria that is shared at lower conservation levels with stalk-forming Betaproteobacteria and dread-forming Zetaproteobacteria. The *sfz* and *sfb* clusters are absent in stalk-less Zetaproteobacteria and Betaproteobacteria, and as previously postulated by Kato et al. (2015), they are likely candidates for encoding the twisted stalk trait. The presence of conserved protein domains for the binding of the signaling messenger c-di-GMP in *sfz* and *sfb* genes together with stalk-former unique genes that putatively encode transcription factors regulating a c-di-GMP dependent signaling cascade for a switch between sessile and motile growth point towards a functional connection between stalk formation and surface attachment. Singer et al. (2011) previously proposed a model according to which stalks serve as holdfasts to anchor cells to surfaces during sessile growth phases. In such a scenario, stalks could provide an advantage over amorphous biofilms as cells can grow within gradients while remaining surface-attached (Chan et al., 2011; Chan et al., 2016). By comparison, stalk-less cells forming amorphous biofilms would have to frequently detach to swarm towards more favorable conditions. Generally, the fact that stalk-formers bear highly conserved unique genes that putatively control cellular motility and surface attachment is supportive of a holdfast function of stalks, though the actual involvement of respective genes in such processes and the extent to which they are linked to stalk formation requires detailed investigation. Our observations reported here require further investigation with lab-based experiments involving differential transcriptomics and gene expression analyses to uncover whether the *sfz* and *sfb* gene clusters indeed encode stalk formation in Zetaproteobacteria and Betaproteobacteria, and to identify whether stalks are functionally linked to surface attachment. In particular, the potential for stalk-less Zetaproteobacteria to harbor alternative stalk formation genes that may be non-operative requires investigation with laboratory experiments.

## 6 Author Contributions

EK, DE, OB and CSC contributed conception and design of the study. EK generated data and performed data analyses. TB conducted SEM sample analyses and supported EK in SEM sample preparation. OB supported EK in workflow design, script writing, and computational data analyses. EK wrote the first draft of the manuscript. All authors contributed to manuscript revision, read and approved the submitted version.

## 7 Acknowledgments

We are grateful to James Byrne for advice on SEM sample preparation and analyses and thank Jake P. Beam and Julia Brown for support in data presentation with R. We also thank Rene Hoover for her comments on a draft of this manuscript. This study was financed by the Office of Naval Research (Grant Nos. N00014-17-1-2641 and N00014-17-1-2640).

