## Supplemental_Information for "Zetaproteobacteria pan-genome reveals candidate gene cluster for twisted stalk biosynthesis and export": Koeksoy_etal_Zetaproteobacteria_Pangenome_SI copy.pdf

### Supplementary Information

**Koeksoy, E., Bezuidt, O.M., Bayer, T., Chan, C.S., Emerson, D.**

### FIGURES

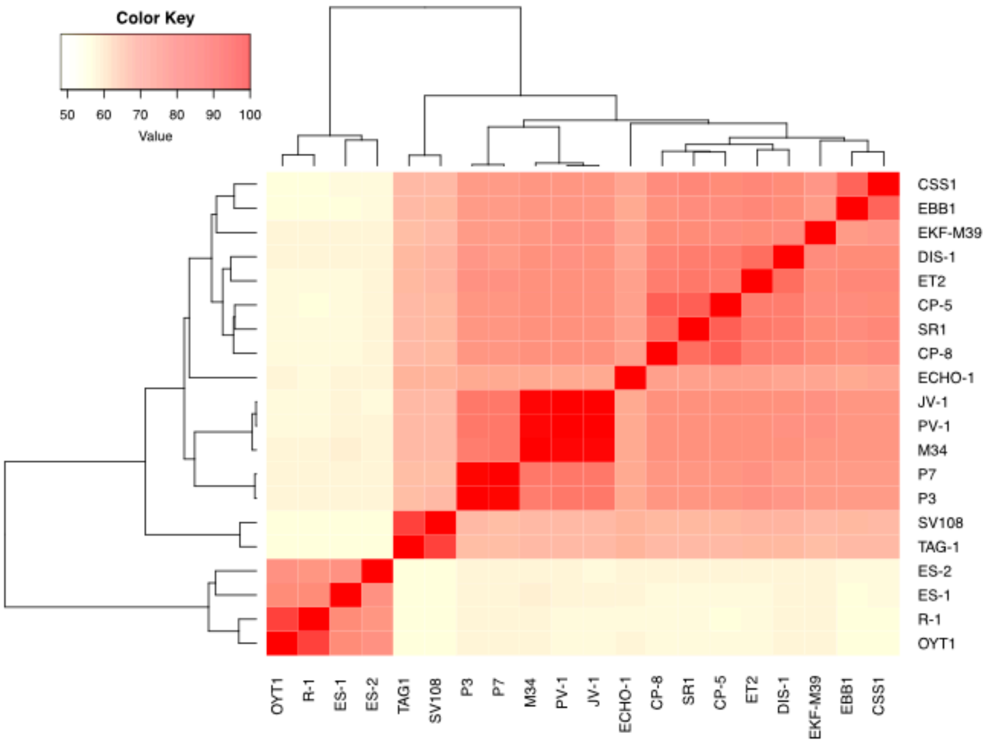


**Figure S1**: Degree of genomic similarity between 16 Zetaproteobacteria and 4 Betaproteobacteria genomes included in this study based on amino acid identities of their protein coding genes. Genomes belonging to the same species are represented in intense red (>95%), whereas lighter colors indicate genomes to fall below the species classification cutoff.

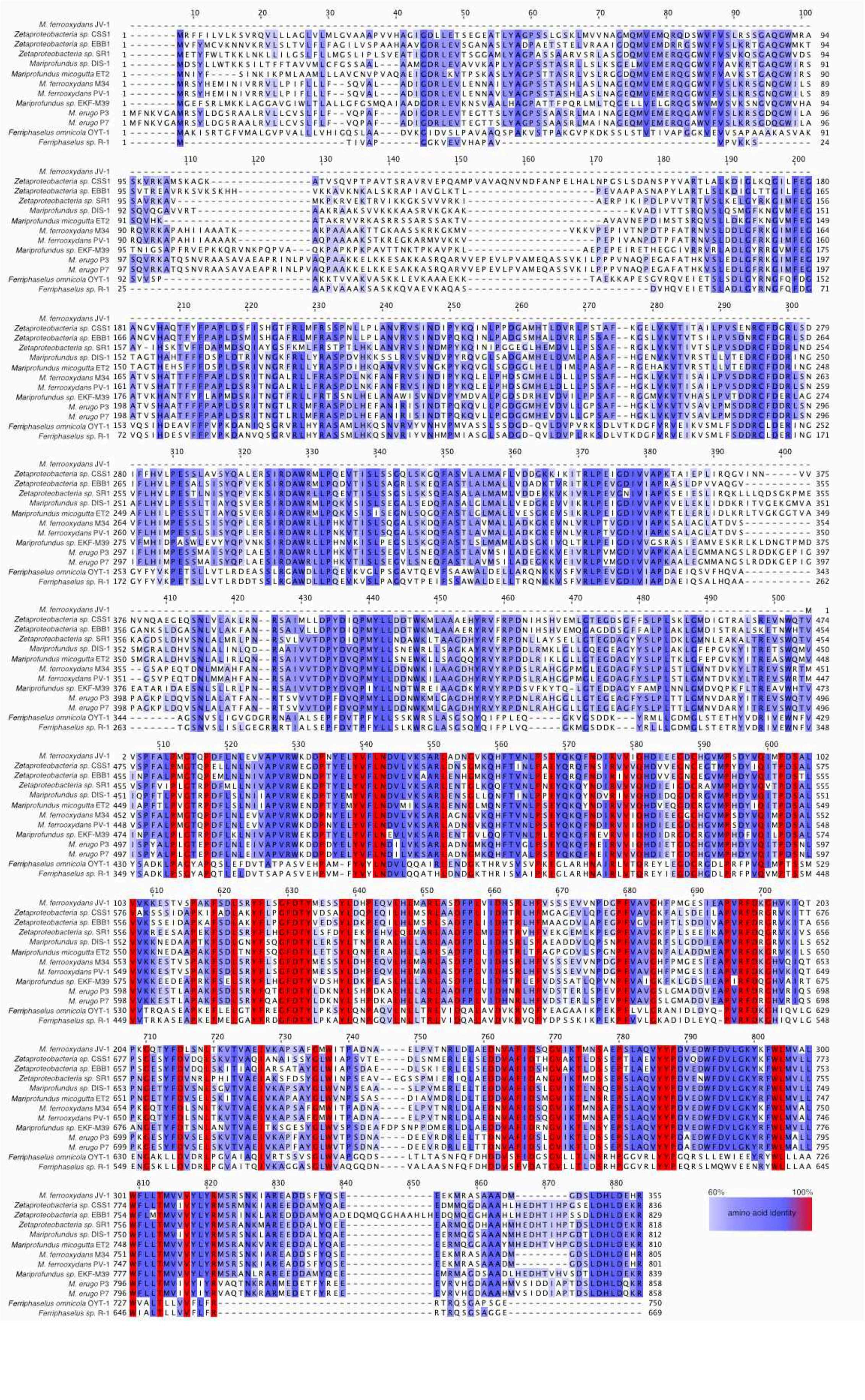
M. ferrooxydans JV-1
Zetaproteobacteria sp. CSS1
Zetaproteobacteria sp. EBB1
Zetaproteobacteria sp. SR1
Mariprofundus sp. DIS-1
Mariprofundus micogutta ET2
M. ferrooxydans M34
M. ferrooxydans PV-1
Mariprofundus sp. EKF-M39
M. erugo P3
M. erugo P7
Ferriphaselus omnicola OYT-1
Ferriphaselus sp. R-1
60%
100%
amino acid identity

**Figure S2:** Alignment of *sfz1 and sfb1* protein sequences in stalk-formers. 100% identical residues are highlighted in red, less conserved residues are shown above 60% amino acid identity.

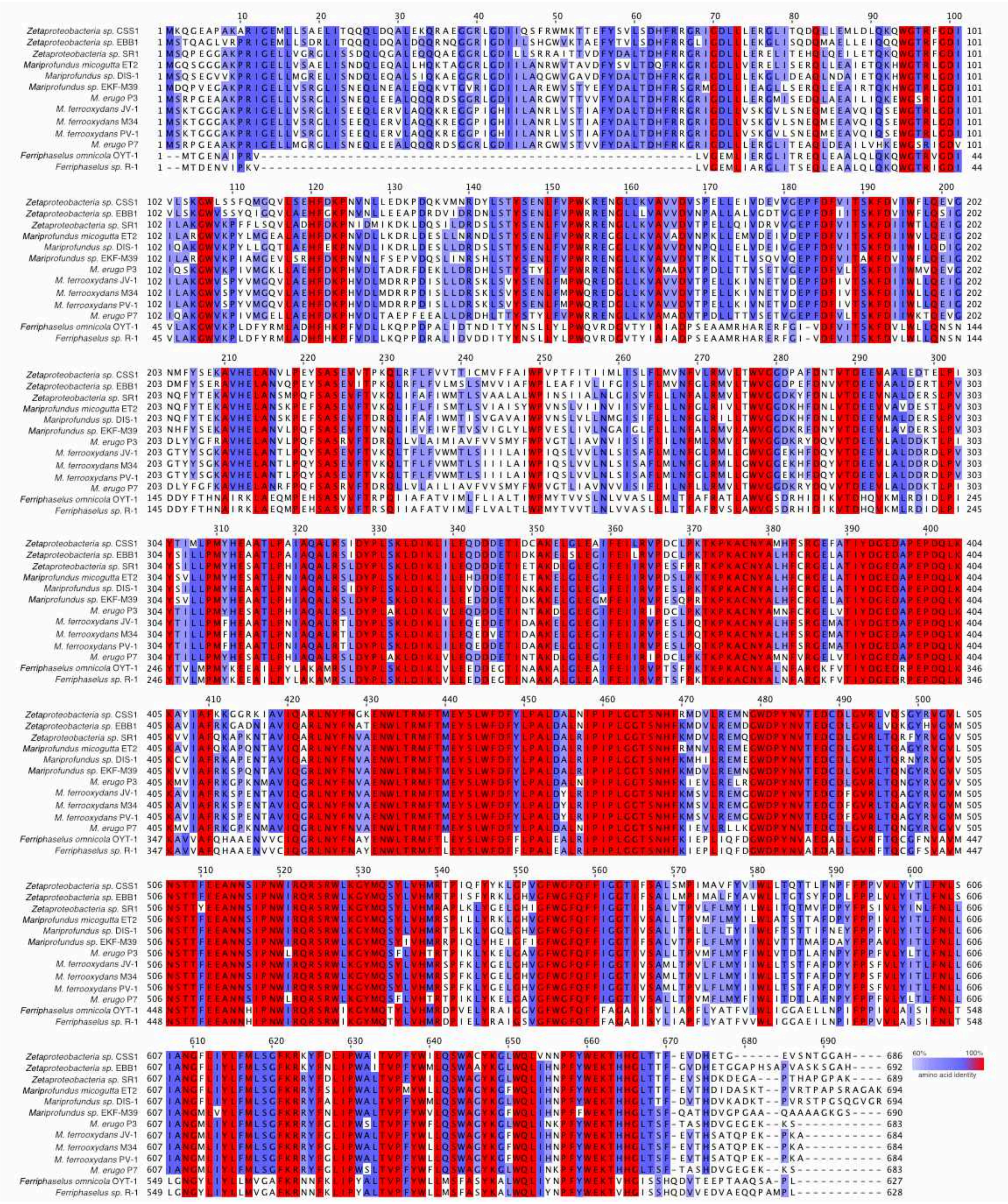

**Figure S3:** Alignment of *sfz2 and sfb2* protein sequences in stalk-formers. 100% identical residues are highlighted in red, less conserved residues are shown above 60% amino acid identity.

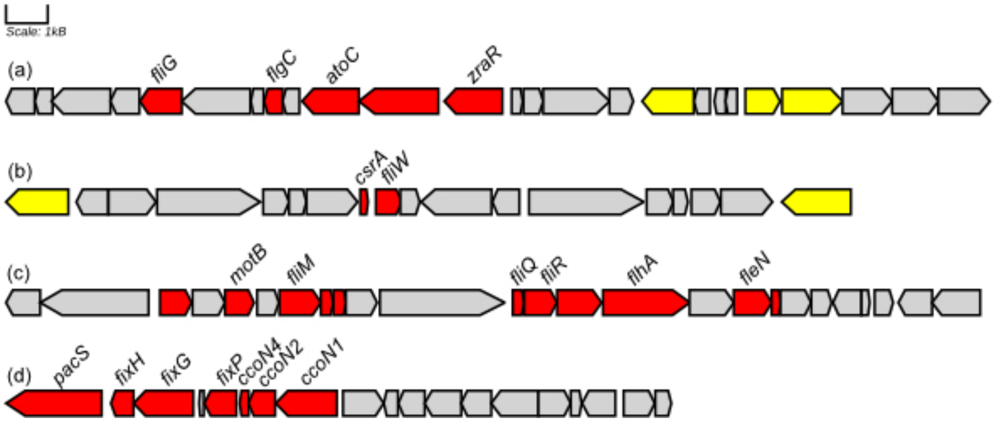


**Fig. S4:** Selected stalk-former unique genes in *M. ferrooxydans* PV-1 (red) with genes that are not unique to stalk-formers (grey) and genes with a potential function in c-di-GMP signaling (highlighted in yellow).

### Supplementary Information - TABLES

**Table S1:** Relative proportions of core, accessory, and strain-specific genes in Zetaproteobacteria pan-genomes computed between 30-90% minimal blastp identity.

| | Minimal blastp identity | | | | | | |
|---|---|---|---|---|---|---|---|
| | 30% | 40% | 50% | 60% | 70% | 80% | 90% |
| **Core** | 10.2 | 10.3 | 8.5 | 5.4 | 2.4 | 0.7 | 0.04 |
| **Accessory** | 48.0 | 46.5 | 44.4 | 42.8 | 38.9 | 31.4 | 22.2 |
| **Strain-specific** | 41.8 | 43.2 | 47.1 | 51.8 | 58.7 | 67.9 | 77.7 |

**Table S2:** Proportions of core, strain-specific, and accessory genes in each isolate genome extracted from the Zetaproteobacteria pan-genome computed at 50% min. blastp identity. Shown values are percentages of total gene numbers.

| Genome | Source environment | Core | Accessory | Strain-specific |
|---|---|---|---|---|
| *M. ferrooxydans* JV-1 | deep sea | 30.8 | 66.7 | 2.5 |
| *M. ferrooxydans* PV-1 | deep sea | 30.6 | 66.3 | 3.1 |
| *M. ferrinatatus* CP8 | estuarine bay | 38.0 | 55.3 | 6.7 |
| *M. erugo* P3 | steel coupon | 32.6 | 60.5 | 7.0 |
| *M. ferrooxydans* M34 | deep sea | 32.3 | 60.2 | 7.4 |
| *Zetaproteobacteria* sp. SR1 | worm burrow | 36.3 | 54.6 | 9.1 |
| *M. erugo* P7 | steel coupon | 31.5 | 59.1 | 9.3 |
| *M. aestuarium* CP5 | estuarine bay | 35.1 | 54.4 | 10.4 |
| *M. micogutta* ET2 | coastal bay | 35.2 | 54.0 | 10.8 |
| *G. bivora* TAG-1 | deep sea | 39.0 | 47.4 | 13.6 |
| *G. bivora* SV-108 | deep sea | 38.5 | 45.0 | 16.5 |
| *Zetaproteobacteria* sp. EBB1 | FeS coupon | 31.9 | 51.4 | 16.7 |
| *Mariprofundus* sp. EKF-M39 | deep sea | 32.0 | 51.0 | 16.9 |
| *Mariprofundus* sp. DIS-1 | coastal bay | 29.6 | 52.7 | 17.6 |
| *Zetaproteobacteria* sp. ECHO1 | worm burrow | 44.2 | 36.3 | 19.4 |
| *Zetaproteobacteria* sp. CSS1 | worm burrow | 31.5 | 48.3 | 20.2 |

**Table S3:** Stalk-former unique genes extracted from Zetaproteobacteria pan-genomes computed at minimal blastp identities between 30-80%. Rows representing *sfz1-6* are shaded in light grey. Genes with potential functional links to a switching response between sessile and motile growth are marked with *.

| Min. blastp identity [%] | Gene | Annotation (prokka) | Min. length [bp] | Max. length [bp] | Average length [bp] |
|---|---|---|---|---|---|
| 80 | *ccoN1* | Cbb3-type cytochrome c oxidase subunit CcoN1 | 1418 | 1466 | 1439 |
| | *flgC** | Flagellar basal-body rod protein FlgC | 446 | 449 | 446 |
| | *flgG** | Flagellar basal-body rod protein FlgG | 551 | 788 | 765 |
| | *flhA** | Flagellar biosynthesis protein FlhA | 2081 | 2087 | 2084 |
| | *fliG** | Flagellar motor switch protein FliG | 1001 | 1019 | 1007 |
| | *fliN** | Flagellar motor switch protein FliN | 296 | 320 | 302 |
| | *fliQ** | hypothetical protein | 224 | 353 | 272 |
| | *csrA** | Carbon storage regulator | 254 | 272 | 259 |
| | HP_80_1 | hypothetical protein | 149 | 206 | 174 |
| | HP_80_2 | hypothetical protein | 641 | 710 | 653 |
| | HP_80_3 | hypothetical protein | 746 | 755 | 749 |
| | *pomA** | Chemotaxis protein PomA | 758 | 758 | 758 |
| | *Ppa* | Inorganic pyrophosphatase | 599 | 635 | 618 |
| | *zraR** | Transcriptional regulatory protein ZraR | 1388 | 1397 | 1392 |
| 70 | *atoC** | Regulatory protein AtoC | 1355 | 1367 | 1361 |
| | *cheB** | Chemotaxis response regulator protein-glutamate methylesterase | 2189 | 2261 | 2208 |
| | *flgE** | Flagellar hook protein FlgE | 1361 | 1730 | 1482 |
| | *flgH** | Flagellar L-ring protein | 695 | 701 | 695 |
| | *flhB_1** | Flagellar biosynthetic protein FlhB | 1067 | 1070 | 1067 |
| | *flhB_2** | Flagellar biosynthetic protein FlhB | 275 | 308 | 286 |
| | *fliF** | hypothetical protein | 1637 | 1658 | 1645 |
| | *fliI** | Flagellum-specific ATP synthase | 1361 | 1454 | 1405 |
| | *fliR** | hypothetical protein | 779 | 782 | 781 |
| | **HP_70_1** | **hypothetical protein (*sfz2*)** | **2051** | **2084** | **2064** |
| | HP_70_2 | hypothetical protein | 398 | 410 | 400 |
| | HP_70_3 | hypothetical protein | 1577 | 1619 | 1589 |
| | HP_70_4 | hypothetical protein | 224 | 323 | 300 |
| | HP_70_5 | hypothetical protein | 566 | 602 | 575 |
| | *motB* | Motility protein B | 698 | 701 | 698 |
| | *ylxH* | Iron-sulfur cluster carrier protein | 857 | 956 | 901 |
| 60 | *fixP* | Cbb3-type cytochrome c oxidase subunit FixP | 851 | 869 | 862 |
| | *flgB** | Flagellar basal body rod protein FlgB | 401 | 407 | 402 |
| | *flgG** | Flagellar basal-body rod protein FlgG | 740 | 743 | 742 |
| | *flgK** | Flagellar hook-associated protein 1 | 1727 | 1745 | 1731 |
| | *fliM** | Flagellar motor switch protein FliM | 986 | 992 | 989 |
| | *fliW** | Flagellar assembly factor FliW | 407 | 458 | 438 |
| | **HP_60_1** | **hypothetical protein (*sfz1*)** | **1067** | **2576** | **2353** |
| | HP_60_2 | hypothetical protein | 842 | 1097 | 924 |
| | HP_60_3 | hypothetical protein | 569 | 596 | 579 |
| | HP_60_4 | hypothetical protein | 1400 | 1427 | 1414 |
| | HP_60_5 | hypothetical protein | 212 | 248 | 220 |
| | **HP_60_6** | **hypothetical protein (*sfz3*)** | **1499** | **1535** | **1509** |

| | | | | | |
|---|---|---|---|---|---|
| | HP_60_7 | hypothetical protein | 482 | 497 | 485 |
| | HP_60_8 | hypothetical protein | 626 | 644 | 636 |
| | HP_60_9 | hypothetical protein | 1277 | 1304 | 1285 |
| | HP_60_10 | hypothetical protein | 380 | 407 | 389 |
| | HP_60_11 | hypothetical protein | 476 | 551 | 505 |
| | *pal* | hypothetical protein | 620 | 758 | 656 |
| | *rcsC* | Sensor histidine kinase RcsC | 1079 | 2114 | 2007 |
| 50 | *flgD** | Basal-body rod modification protein FlgD | 665 | 683 | 671 |
| | **HP_50_1** | **hypothetical protein (*sfz6*)** | **3311** | **7352** | **4899** |
| | HP_50_2 | hypothetical protein | 695 | 737 | 713 |
| | HP_50_3 | hypothetical protein | 686 | 785 | 720 |
| | HP_50_4 | hypothetical protein | 278 | 344 | 318 |
| | HP_50_5 | hypothetical protein | 1520 | 1541 | 1530 |
| | HP_50_6 | hypothetical protein | 1907 | 1985 | 1952 |
| | HP_50_7 | hypothetical protein | 254 | 263 | 257 |
| | HP_50_8 | hypothetical protein | 380 | 404 | 383 |
| | HP_50_9 | hypothetical protein | 641 | 674 | 655 |
| | HP_50_10 | hypothetical protein | 212 | 230 | 219 |
| | HP_50_11 | hypothetical protein | 440 | 518 | 482 |
| | **HP_50_12** | **hypothetical protein (*sfz4*)** | **971** | **1037** | **1006** |
| | **HP_50_13** | **hypothetical protein (*sfz5*)** | **929** | **983** | **946** |
| | HP_50_14 | hypothetical protein | 281 | 350 | 301 |
| | HP_50_15 | hypothetical protein | 1406 | 1439 | 1430 |
| | *pacS* | putative copper-transporting ATPase PacS | 2291 | 2591 | 2423 |
| | *pckA* | Phosphoenolpyruvate carboxykinase (ATP) | 995 | 1595 | 1532 |
| 40 | HP_40_1 | hypothetical protein | 701 | 1007 | 759 |
| | HP_40_2 | hypothetical protein | 437 | 452 | 444 |
| | HP_40_3 | hypothetical protein | 458 | 494 | 464 |
| | HP_40_4 | hypothetical protein | 1619 | 2213 | 1763 |
| | HP_40_5 | hypothetical protein | 497 | 509 | 500 |
| | HP_40_6 | hypothetical protein | 1436 | 1472 | 1444 |
| | HP_40_7 | hypothetical protein | 293 | 398 | 318 |
| | HP_40_8 | hypothetical protein | 623 | 809 | 716 |
| | HP_40_9 | hypothetical protein | 1010 | 1526 | 1238 |
| 30 | HP_30_1 | hypothetical protein | 809 | 818 | 812 |
| | HP_30_2 | hypothetical protein | 1229 | 1253 | 1239 |
| | HP_30_3 | hypothetical protein | 1691 | 2018 | 1760 |
| | HP_30_4 | hypothetical protein | 278 | 323 | 291 |
| | HP_30_5 | hypothetical protein | 482 | 575 | 530 |
| | *rcsC* | Sensor histidine kinase RcsC | 401 | 2738 | 1940 |

**Table S4:** Top 10 COGs in strain-specific genes of the Zetaproteobacteria pan-genome computed at 50% minimal blastp identity.

| COG category | COG functional annotation | Counts |
|---|---|---|
| COG3706 | Response regulator containing a CheY-like receiver domain and a GGDEF domain | 246 |
| COG1196 | Chromosome segregation ATPases | 177 |
| COG2197 | Response regulator containing a CheY-like receiver domain and an HTH DNA-binding domain | 175 |
| COG2204 | Response regulator containing CheY-like receiver, AAA-type ATPase, and DNA-binding domains | 172 |
| COG3437 | Response regulator containing a CheY-like receiver domain and an HD-GYP domain | 171 |
| COG0784 | FOG: CheY-like receiver | 169 |
| COG0745 | Response regulators consisting of a CheY-like receiver domain and a winged-helix DNA-binding domain | 166 |
| COG4753 | Response regulator containing CheY-like receiver domain and AraC-type DNA-binding domain | 165 |
| COG2202 | FOG: PAS/PAC domain | 156 |
| COG4566 | Response regulator | 148 |
